# A DNA Methylation Fingerprint of Cellular Senescence

**DOI:** 10.1101/674580

**Authors:** Morgan E. Levine, Diana Leung, Christopher Minteer, John Gonzalez

**Affiliations:** Department of Pathology, Yale School of Medicine, New Haven, CT, USA, 06519

## Abstract

Recent years have witnessed an influx of therapies aimed at targeting the biological aging process. While many hold promise for potentially delaying the simultaneous onset of multiple chronic conditions, a major hurdle in developing interventions to slow human aging is the lack of reliable and valid endpoints from which to evaluate potential candidates. The aim of this study was to develop a biomarker that could serve as a potential endophenotype in evaluation of one of the most promising and exciting aging therapeutics being developed—senolytics. Senolytics are compounds which selectively clear senescent cells by targeting anti-apoptotic pathways. Using DNAm data from fibroblasts and mesenchymal stromal cells (MSC) in culture, we developed a predictor of cellular senescence that related to three distinct senescence inducers (replicative, oncogene induced (OIS), and ionizing radiation (IR)). Our measure, termed DNAmSen, showed expected classification of validation data from embryonic stem cells (ESC), induced pluripotent stem cells (iPSC), and near senescent cells in culture. Further, using bulk data from whole blood, peripheral tissue samples, and postmortem tissue, we observed robust correlations between age and DNAmSen. We also observed age adjusted associations between DNAmSen and idiopathic pulmonary fibrosis (IPS), COPD, lung cancer, and Werner syndrome compared to controls. When characterizing the 88 CpGs in the DNAmSen measure, we observed a enrichment for those located in enhancer regions and regulatory regions marked by DNase I hypersensitive sites (DHSs). Interestingly most of the CpGs were observed to be moderately hypomethyled in early passage cells, yet exhibited further hypomethylation upon induction of cellular senescence, suggesting senescence is accompanied by programmatic alterations to the epigenome, rather than entropic drift.

## INTRODUCTION

At a population level, incidence rates for the majority of chronic diseases, disability, and mortality increase exponentially as a function of age^1^. Nevertheless, significant between-person variance is observed in age-adjusted risks for most of these conditions. This stems from the simple fact that while chronological aging increases at a fixed rate for everyone, biological aging—the rate at which the body moves from young/vibrant to old/sick—is variable and is hypothesized to play a causal role in accelerating or decelerating disease pathogenesis^2–4^. As such, identifying the underlying mechanisms that control the rate of biological aging, and developing novel therapeutics to target them will produce enormous personal, societal, and economic impact, by reducing diseases burden and extending not just lifespan, but also healthspan.

Coincidentally, studies in model organisms have demonstrated that biological aging is malleable, either through genetic and/or environmental manipulation^5^. However, translational applications to slow and/or reverse biological aging in humans has thus far remained elusive. Despite this, there is growing consensus in the field when it comes to defining the most prominent cellular and molecular hallmarks of aging from which to develop potential therapeutic targets^6^. One such hallmark is cellular senescence. Cellular senescence is a cellular state defined by proliferative arrest^7,8^. It is often a characteristic of genomic instability or cellular stress, and can be induced via telomere attrition, DNA damage, mutations in oncogenes, inflammation, elevated glucose, irradiation, and/or mitochondrial dysfunction. Senescent cell accumulation in aging tissues is hypothesized to play a role in the etiology of various age-related conditions, as a result of its corresponding transcriptional activation, termed the senescence-associated secretory phenotype (SASP)^9^. Although they remain non-proliferative, senescent cells have been shown to actively secrete cytokines, chemokines, proteases, and other damaging agents, thus contributing to a chronic proinflammatory environment^10,11^.

In recent years, there has been a growing movement to develop therapeutics to facilitate clearance of these toxic cells^8^. The first compounds to selectively clear senescent cells—termed senolytics—were discovered in 2015^12–14^. These were based on targeting of the pro-survival/anti-apoptotic pathways that were found to be upregulated in senescence. To date, a number of senolytic drugs have been developed, or are in the development pipeline^8^. However, while morbidity/functioning traits have typically been used to demonstrate efficacy, a major hurdle for developing senolytics for human trials is the lack of valid and reliable biomarkers^15,16^. Up until now, a number of potential assays have been used to infer senescence, including tissue staining for β-galactosidase positive (SA-βgal^+^) cells, p16^INK4A+^ expression, SASP factor activation, and damage foci (e.g. γH2.AX). However, none of these biomarkers is specific enough to be used alone for clinical evaluation of senolytic therapies^8^.

Interestingly, it has been well established that senescence is also marked by another hallmark of aging—epigenetic alterations—which includes changes in DNA methylation (DNAm)^17–19^. Epigenetic alterations are believed to play an important role in establishment and maintenance of senescence by silencing proliferation-promoting genes. Recent work using Human BJ Fibroblasts^17^ has shown that replicative senescence (RS) exhibits an almost programmatic change in DNAm, characterized by hypermethylation of promoter regions for genes involved in biosynthetic and metabolic processes. Interestingly, CpG Island promoter hypermethylation, and global hypomethylation are observed in senescence, normal aging, and cancer. Nevertheless, many studies examine only a single type of senescence—typically RS—making it difficult to differentiate DNAm signals that reflect the type of inducer, versus senescence itself. Further, even among the studies investigating various types of senescence, most have pointed to the contrasting DNAm patterns, rather than identifying a universal signature.

Previously, we and others have employed DNAm data to develop biomarkers that universally capture normal aging across diverse tissues and cell types^20–24^. These ‘epigenetic ages’ can be contrasted against the chronological age of the sample donor to differentiate accelerated versus decelerated aging. Furthermore, we have shown that the difference in epigenetic and chronological age is a robust predictor of morbidity and mortality risk^22^. Nevertheless, the underlying biology of what these measures are capturing is unclear, decreasing their potential application for clinical trials^25^.

As a result, the aim of this paper is to bridge these two approaches by utilize DNAm data from in vitro experiments to generate a novel epigenetic biomarker that specifically captures cellular senescence in human samples. In doing so we consider three types of senescence—replicative senescence (RS), oncogene induced senescence (OIS), and Ionizing Radiation-Induced (IRIS) senescence—in two cell types, BJ Fibroblasts and Mesenchymal Stromal Cells (MSC).

## RESULTS

### Development of a Senescence DNAm Biomarker

The primary data used to train a DNAm biomarker of senescence came from two studies, one with senescence in Human BJ Fibroblasts and the other with senescence in Mesenchymal Stromal Cells (MSC) (Figure 1).

**Figure 1:**
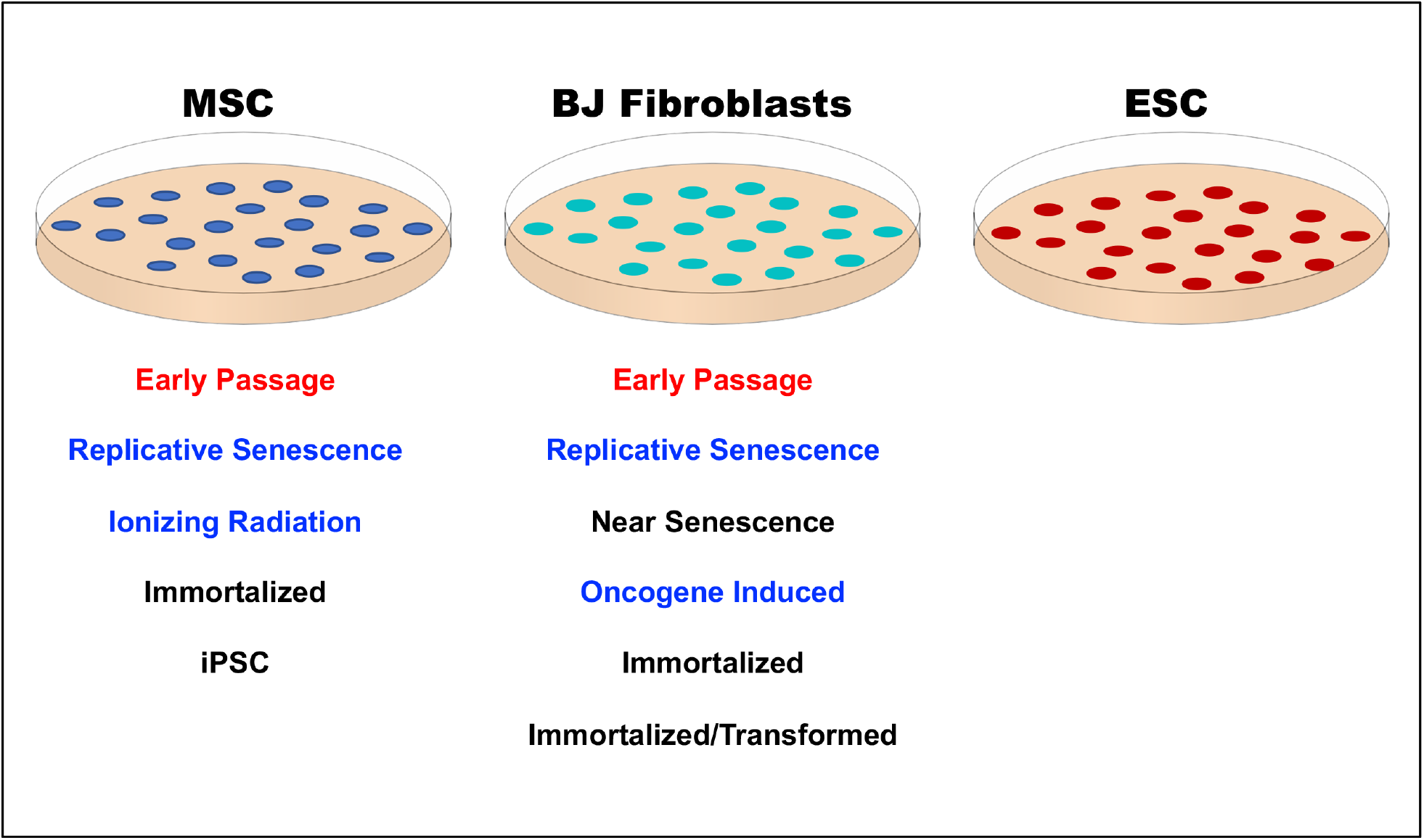
Framework for development of DNAmSen classifier. Data from mesenchymal stem cells, BJ fibroblasts, and embryonic stem cells was used to train and validate a DNAm based predictor of cellular senescence. For training DNAm from early passage cells (red) were compared to senescent cells (blue), considering three types of inducers.

Prior to pooling MSC and Fibroblast samples, we used t-Distributed Stochastic Neighbor Embedding (t-SNE) to compare the signals between the two samples to determine if we needed to correct for batch effects before applying the supervised machine learning approach. As shown in Figure 2A, MSC and Fibroblasts clustered separately for component 2. However, upon further inspection this distinction appears to have a biological rather than a technical basis. For instance, it also fully distinguishes ESC and iPSC from MSC and Fibroblasts. Even within the Fibroblast samples, there is a clear clustering by condition, with EP cells having the highest (closest to EP) values for component 2.

**Figure 2:**
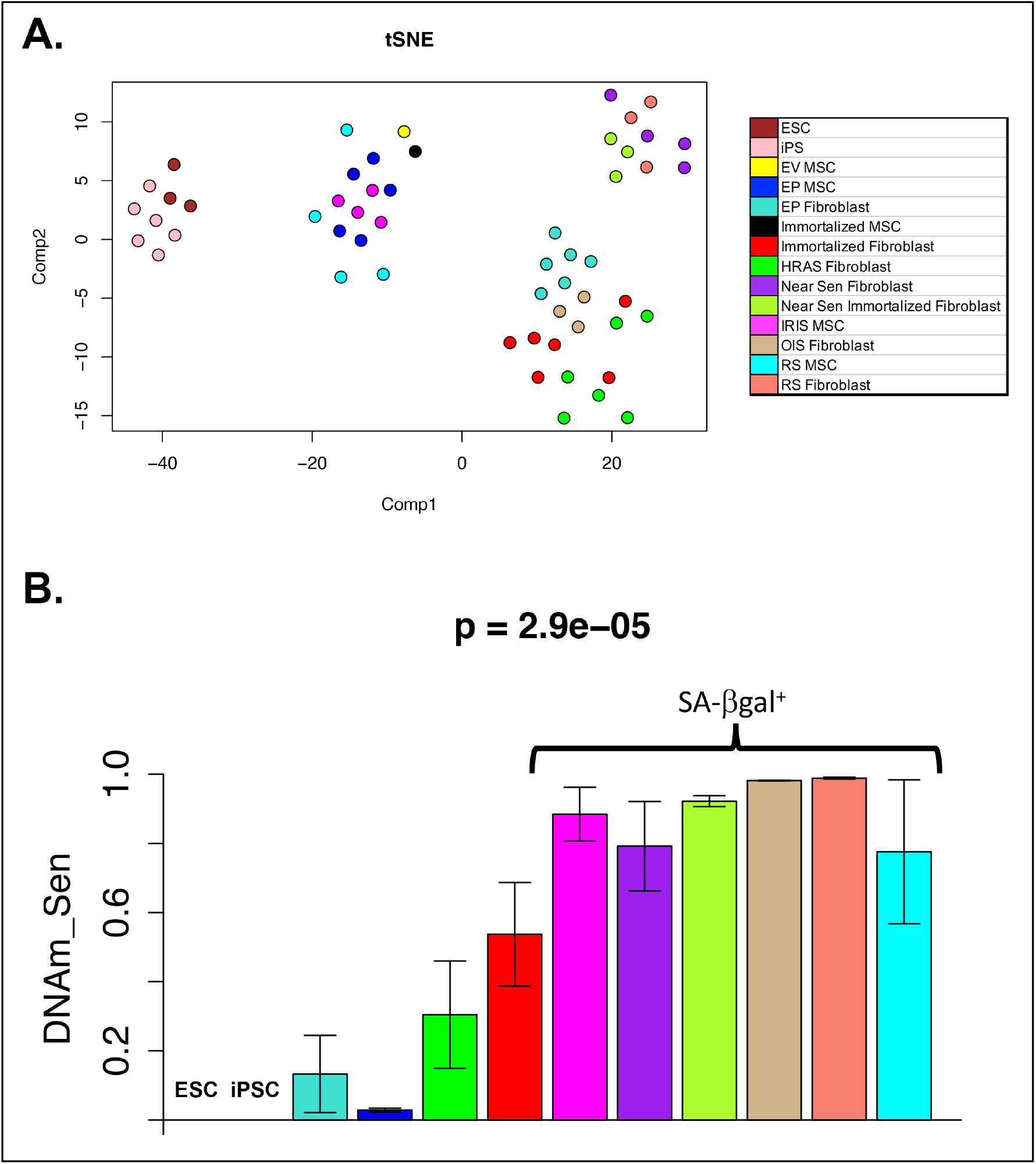
Distinct DNAm profiles of senescent and non-senescent cells. t-Distributed Stochastic Neighbor Embedding (t-SNE) was applied in order to cluster cell types based on methylation patterns across more than 450,000 CpG cites (2A). Cells were differentiated based on both components suggesting that they exhibit unique DNAm patterns. After training a supervised classier of senescence, the resulting equation was applied and we compared the predictor values across cell types (2B). We observed much higher levels for all SA-βgal+ cells, while the lowest levels were observed for ESC and iPSC.

Based on this, data from cultured MSC and Fibroblasts were pooled, and a supervised machine learning approach (elastic net regression) was used to generate a classifier of senescence (Figure 1). To do so, we compared the three types of senescence—totaling 12 senescence samples to the MSC and fibroblast EP samples (n=10). This resulted in a predictor based on 88 CpGs. Figure 2B shows the levels for the resulting DNAm senescence biomarker by condition. While the OIS, RS, IRIS, and EP were used as training data, we still find that ESC and iPSC are classified as completely non-senescent (DNAmSen=0). Conversely, the near senescent cells, which stain positive for SA-βgal reach almost the same level as the senescent cells (DNAmSen>0.8). Immortalized and immortalized/transformed cells have a slight increase (DNAmSen of about 0.3-0.5), but are still significantly distinct from the senescent cells.

### In Vivo Age Validation

The utility of a biomarker for senescence will be based on its performance in readily available samples, such as whole blood. As a result, we calculated DNAmSen using the DNAm data from whole blood for persons ages 19 to 101 years (GSE40279). We found that DNAmSen was correlated with age at r=0.52, p=2.6e-64 (Figure 3A). Moreover, when examining DNAmSen by age group, we observe a very clear separation in levels between persons <40, 50-65, 85+ years—compared to those less that 40 years old, those 50-65 were just over one-half standard deviation higher; while those 85+ were about one and a half standard deviations higher (Figure 3B).

**Figure 3:**
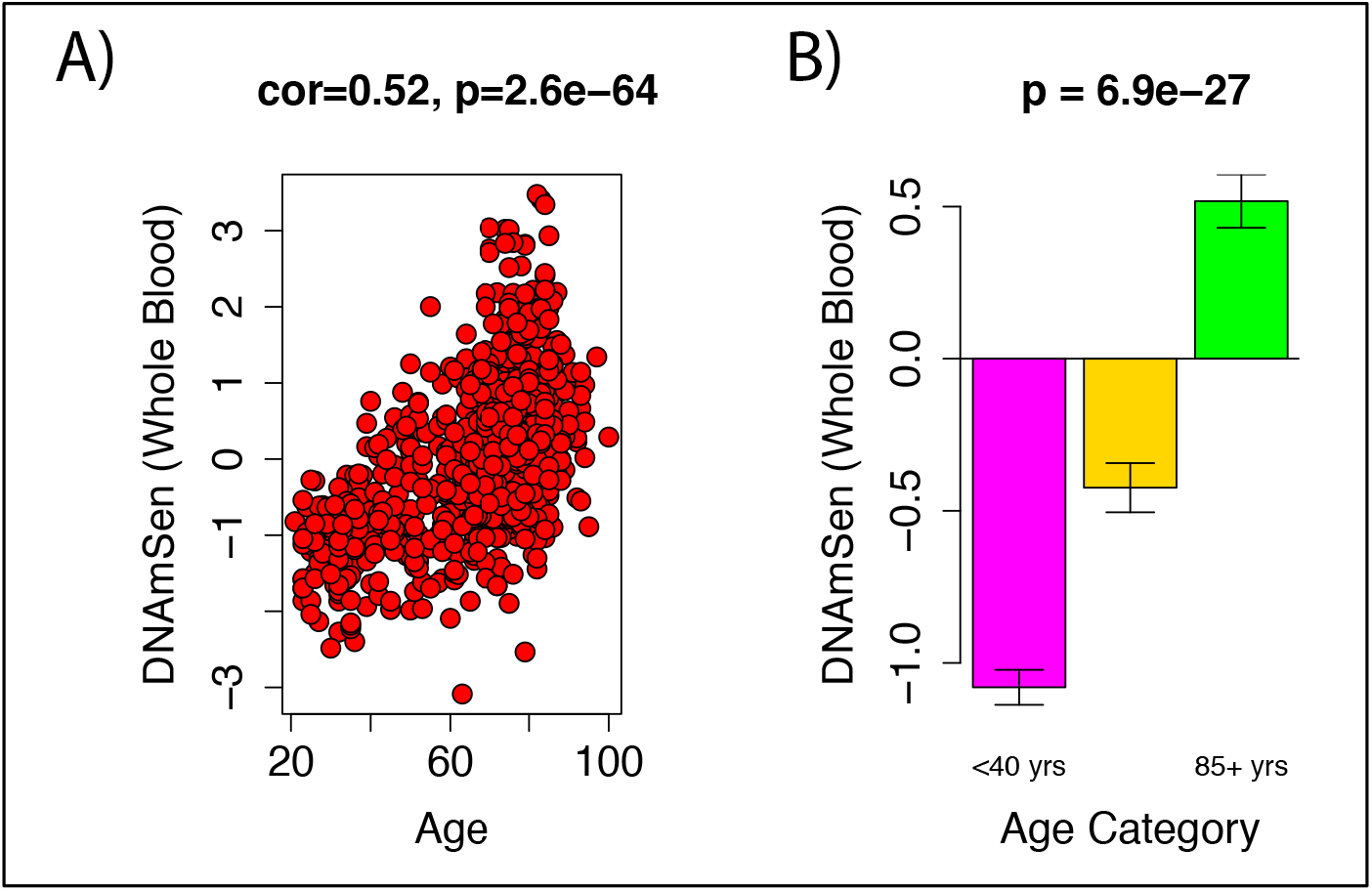
Age differences in DNAmSen estimated in whole blood. The equation developed from cell culture samples was applied to bulk samples from whole blood. When evaluating the age correlation, we find that DNAmSen was strongly/moderately correlated with chronological age in whole blood at r=0.52 (3A). When evaluating this in age-startified groups, we find that younger adults (20-39), middle-aged adults (50-65), and older adults (85+) display distinctly different leves of DNAmSen in whole blood.

Next, we also examined the age correlations in epidermal and dermal skin samples (GSE51954), in which we find robust age correlation in both (Figure 4A and 4B).

**Figure 4:**
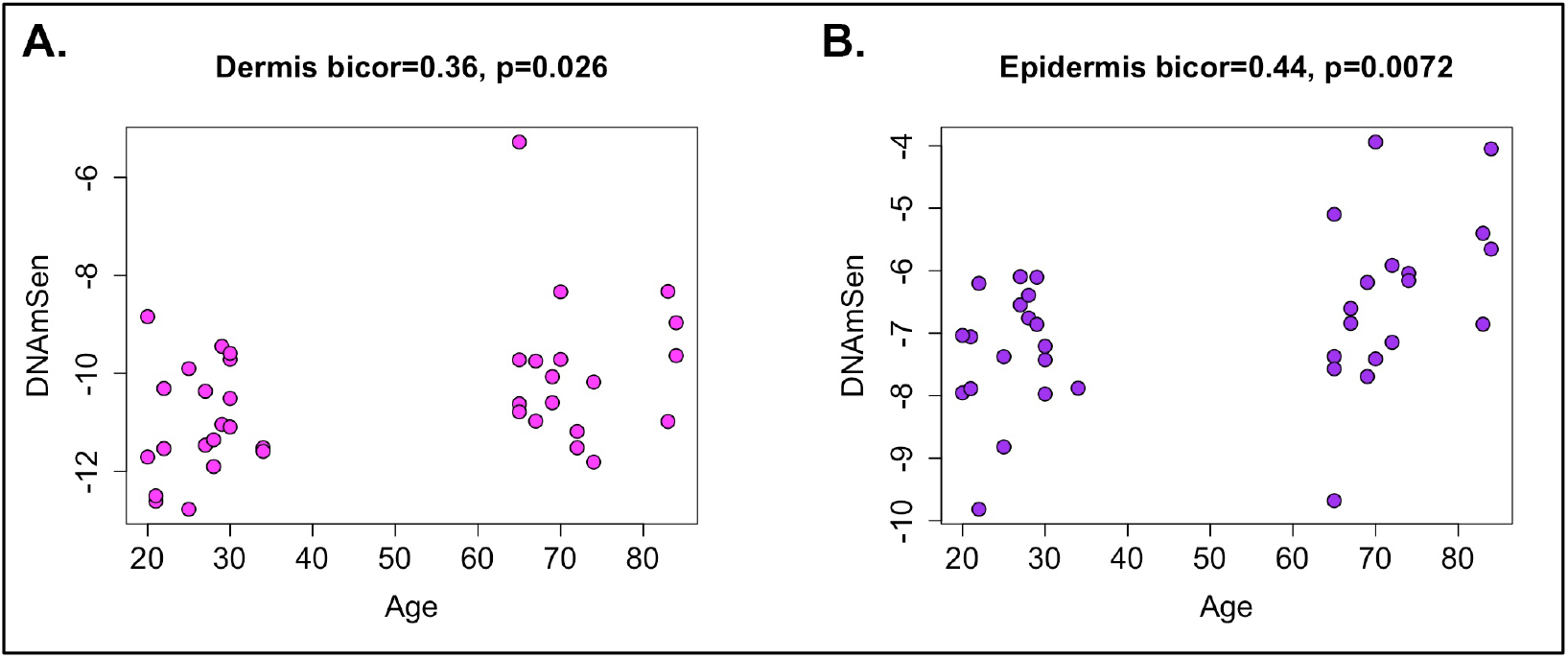
Age correlations for DNAmSen in skin samples from dermis and epidermis. We estimated DNAmSen in skin samples from both dermis and epidermis and found significant age correlations in both: dermis (r=0.36), epidermis (r=0.44).

Finally, we also examine changes in DNAmSen in various tissues during fetal development (Figure 5). Overall, we find that amnion, kidney, skin, eye, muscle, and heart all show robust increases in DNAmSen with increasing gestational age. For instance, amnion—the membrane which encloses the fetus—exhibits a correlation of r=0.95 (p=3.0e-4) between gestational age and DNAmSen. Heart and skin were also found to have robust correlations of r=0.99, and r=0.92, respectively; yet, further validation should be done given that only 4-5 samples were available for each.

**Figure 5:**
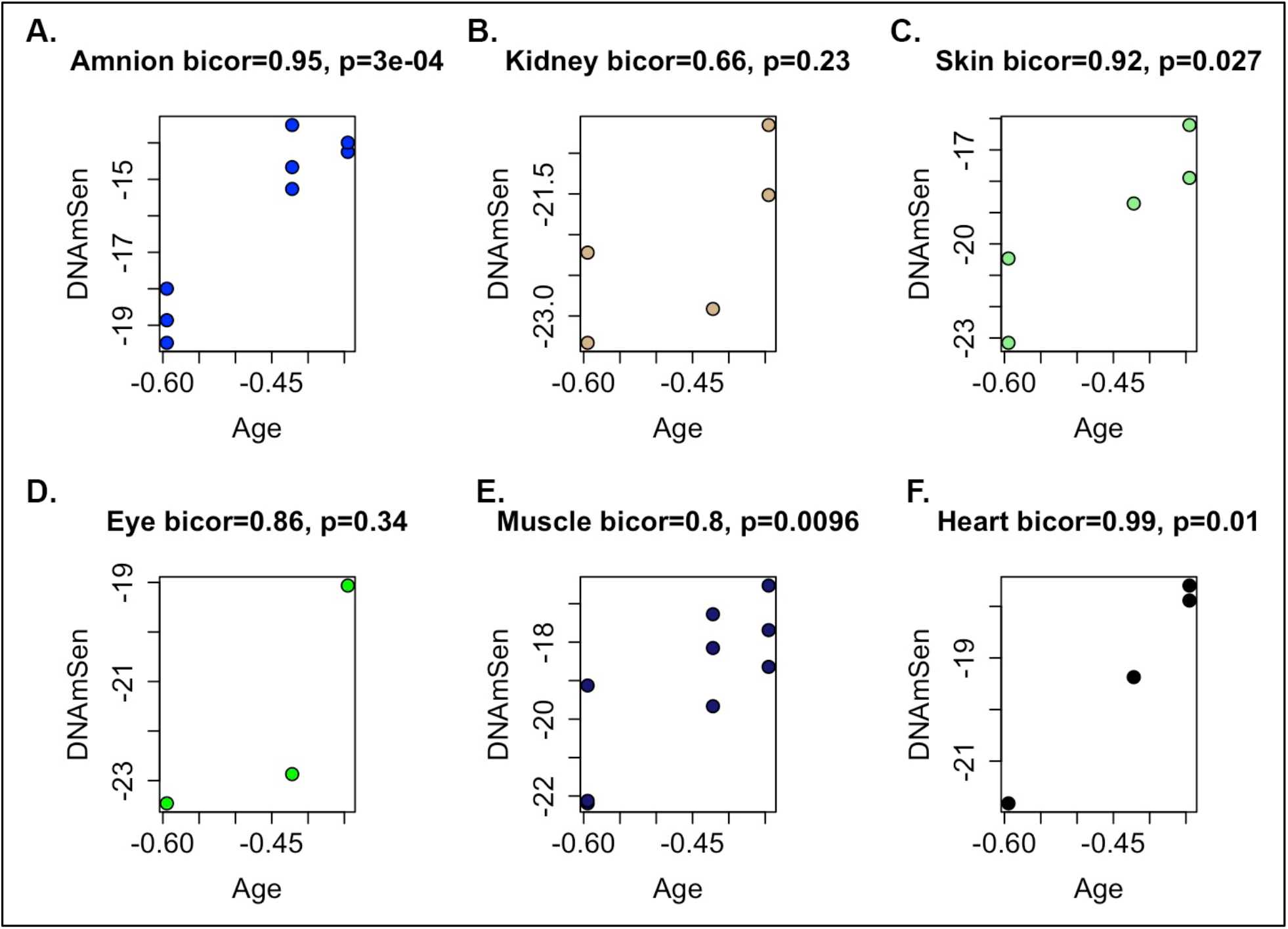
Gestational age correlations in fetal tissue from postmortem samples. We estimated age correlations for DNAmSen measured in a number of fetal tissue samples. Overall, we found extremely strong correlations between for all tissues, ranging from 0.66 in kidney (5B) to 0.99 in heart (5F).

### In Vivo Validation of Aging-Related Outcomes

DNAm measured in various tissue cell samples for different age-related conditions was used to further validate DNAmSen. For instance, based on data from Heyn et al., 2013 (GSE42865) we examined DNAmSen in two progeria conditions (Table 1)—Werner Syndrome (WS) and Hutchinson-Gilford Progeria (HGP). We compared Immortalized B cells from normal controls to those with *lamin A* (LMNA) mutation (WS), non-mutant WS and HGP patients, and *Werner syndrome RecQ helicase like* (WRN) mutation (WS). Despite very small sample sizes, we observed significantly higher DNAmSen among WRN mutant carriers, such that they had nearly a two standard deviation increase in the measure (β=1.84, p=4.19E-3). However, these results were not age-adjusted, because information on age was not available for these samples, Thus, to account for this, we applied the Horvath pan-tissue DNAmAge clock to estimate chronological age based on DNAm. We find that after includion of Horvath DNAmAge, the associations were maintained (β=1.82, p=6.11E-3).

**Table 1:** Associations between DNAmSen and Progeria Mutations.

|  | Model 1 |  | Model 2 |  |
| --- | --- | --- | --- | --- |
|  | Coefficient | p-value | Coefficient | p-value |
| LMNA mutation | 0.8 | 1.76E-01 | 1.1 | 1.13E-01 |
| Non-mutant patient | -0.37 | 3.27E-01 | -0.18 | 6.51E-01 |
| WRN mutation | 1.84 | 4.19E-03 | 1.82 | 6.11E-03 |
| Horvath Clock |  |  | 0.02 | 3.08E-01 |
| R2 | 0.853 |  | 0.883 |  |

Next, using data from GSE63704, DNAmSen in lungs from persons with COPD, Idiopathic Pulmonary Fibrosis (IPF), and lung tumors was compared to DNAmSen in lungs from health controls. We observed that all three disease conditions exhibited elevated DNAmSen compared to controls (Table 2), such that lung tumors had an almost one standard deviation increase (β=0.91, p=1.22E-3), COPD lungs showed over half a standard deviation increase in DNAmSen (β=0.65, p=4.28E-3), and IPF lungs exhibited about a half standard deviation increase in DNAmSen (β=0.51, p=1.91E-2). As with the progeria data, age information was not available for these samples, thus we used the Horvath DNAmAge clock as a surrogate to adjust for chronological age. When adjusting for this measure, we find that the results for cancer and COPD are maintained (β_cancer_=1.10, p_cancer_=9.89E-5; and β_COPD_=0.53, p_COPD_=1.80E-2), yet the result for IPF becomes non-significant (β=−0.002, p=9.92E-1).

**Table 2:** Associations between DNAmSen and Diseases of the Lung.

|  | Model 1 |  | Model 2 |  |
| --- | --- | --- | --- | --- |
|  | Coefficient | p-value | Coefficient | p-value |
| Cancer | 0.91 | 1.22E-03 | 1.1 | 9.89E-05 |
| COPD | 0.65 | 4.28E-03 | 0.53 | 1.80E-02 |
| Fibrosis | 0.51 | 1.91E-02 | -0.002 | 9.92E-01 |
| Horvath Clock | -- | -- | 0.02 | 3.95E-03 |
| R2 | 0.106 |  | 0.164 |  |

### DNAmSen CpG Characteristics

We examined the characteristics of the 88 CpGs in the DNAmSen score and compared them to all the CpGs shared between the 450k and the EPIC Illumina arrays. Overall, CpGs are distributed over the whole genome and don’t appear to cluster in specific chromosomes or genomic locations (Figure 6A). However, we do find a nearly two-fold enrichment for CpGs located in enhancer regions. For instance, nearly 41% of CpGs in our senescence measure were located in enhancer regions, as identified by ENCODE. Conversely, only 21% of the CpGs on the array are located in ENCODE defined enhancer regions. Based on Fisher’s exact test, this suggests a significant enrichment among our 88 probes, versus what is expected by chance (p=0.0013).

**Figure 6:**
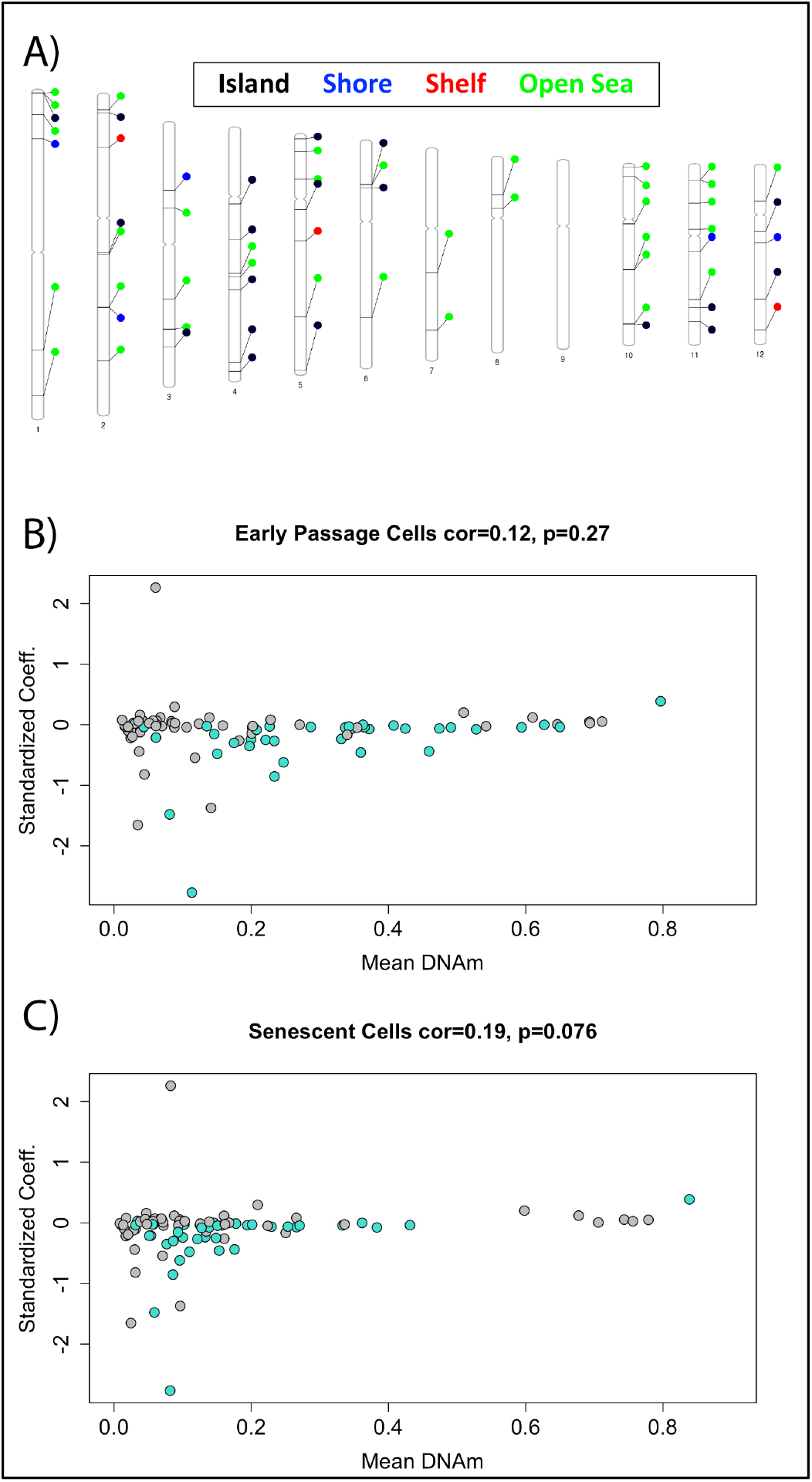
Characteristic of the 88 CpGs in DNAmSen. We found that the 88 CpGs that made up our senescence biomarker were more-or-less evenly distributed across the genome (6A). Interestingly, we observed that the changes likely did not represent epigenetic drift but rather a programmatic change, given that CpGs that were hypomethylated in early passage cells tended to have negative coefficients, suggesting that DNAm was decreased further in senescence. This was supported by the shift to the left in the mean DNAm for senescent cells (6C). For 6B and 6C, CpGs in enhancer regions are indicated in turquoise and show a shift towards more extreme hypomethyltion with senescence.

Similarly, we also observe a significant enrichment for CpGs located in regulatory regions of the DNA, marked by DNase I hypersensitive sites (DHSs). As with the enhancers, we observed nearly two-times the proportion of DHS CpGs for the 88 senescence probes compared to what is observed among all probes on the array (enrichment=1.94, p=0.0089). Finally, there was a slight enrichment for CpGs located in CpG islands (CGI)—just over 30% for the 450k probes, versus 39% for the senescence probes. However, this does not represent a significant enrichment based on Fisher’s exact test (p=0.286). Finally, there was also suggestive enrichment for CpGs that were classified as being promoter associated (enrichment=1.44, p=0.094).

Lastly, we examined whether DNAm differences between senescence and early passage that are captured by our measure represent entropic drift (regression towards the mean). As shown in Figure 6B, generally we find that most CpGs have a coefficients at or near zero (e.g. contribute very little information) to the DNAmSen predictor. However, of those with larger contributions/coefficients, the majority have negative coefficients, suggesting a trend toward hypomethylation in senescence. Yet perhaps more interesting is the observation that the majority of CpGs with robust negative coefficients, are already hypomethylated in early passage cells, yet have DNAm levels that are further reduced in senescence. This suggests that the DNAm changes in senescence are not indicative of epigenetic drift and or entropic alterations (would trend towards 0.5), but rather appear to represent a more programmatic response to senescence inducing stimuli.

## DISCUSSION

Using DNAm data from fibroblasts and mesenchymal stromal cells in culture, we applied supervised machine learning to train a classifier of cellular senescence to serve as a useful biomarker for both clinical trials and basic research. Recently, there has been increasing momentum when it comes to developing therapeutics to target the biological aging process. However, our ability to effectively translate many of these interventions to humans, and in turn comprehensively assess their efficacy in clinical trials, will require valid and reliable endpoints that capture aging and its respective hallmarks^25^.

Biomarkers based on genome-wide DNAm have proven to be reliable indicators of aging processes^20,22–24^. However, many of the existing methylation aging biomarkers—often referred to as epigenetic clocks—are not well understood biologically and likely capture a variety of age-related alterations. Thus, it should be more beneficial to develop aging biomarkers that are meant to specifically capture certain hallmarks that existing therapies aim to target.

One of the most promising aging therapies being developed are senolytics^8^. These interventions are aimed at targeting the anti-apoptotic pathways of senescent cells, facilitating their selective removal. However, one difficulty in evaluating these therapies in human clinical trials centers on the lack of agreed upon measures to quantify the proportions (or change in proportion) of senescent cells. In response, we utilized DNAm data from two distinct cell types, and three distinct senescence inducers to identify an “epigenetic fingerprint” of cellular senescence. Using supervised machine learning, we constructed a classifier of senescence that incorporated information from 88 genome-wide CpGs. Using validation samples, we observed that, as expected, embryonic stem cells and induced pluripotent stem cells were classified as completely non-senescent and even exhibited lower senescent states than early passage foreskin fibroblasts and MSC. Further, fibroblasts that had been passaged until they were nearly senescent—as indicated by SA-βgal^+^ staining coupled with continued passaging capacity—displayed high levels of DNAmSen that were near, yet not quite at the levels of fully senescent cells.

Yet, in order for these measure to be applicable to human clinical trials, they need to produce valid predictions in peripheral samples from living patients. To demonstrate efficacy, we tested the age correlations for DNAmSen using various tissues and cells from both adult and fetal samples. We observed robust age correlations, particularly for samples from adult whole blood and skin, as well as multiple fetal tissues. In addition to age correlations, we also tested whether DNAmSen was higher in various disease conditions, after adjusting for chronological age (or surrogate epigenetic age measures). We observed elevated DNAmSen for lung samples from patients with IPF, COPD, or cancer versus controls; as well as immortalized B cells carriers of WRN mutations versus controls.

When considering the landscape of our DNAmSen measure we observed approximately two-fold enrichment for CpGs located in enhancer regions and regulatory regions, marked by DNase I hypersensitive sites (DHSs). Perhaps more interestingly, we find that there is a trend for DNAm that is captured in our measure to go from already hypomethylated in early passage cells to more extreme levels of hypomethylation with induction of cellular senescence. Assuming the changes in DNAm represented epigenetic drift (entropic processes) we would have expected DNAm of hypomethylated regions to increase, towards mean levels of around 0.50. However, the shift from already low levels of DNAm to even more extreme absence of DNAm suggests that this is a highly regulated alteration, rather than random error.

Overall, we were able to identify a DNAm based measure of cellular senescence that is not unique to a single cell type nor a single inducer (e.g. replicative). At the same time, this measure was able to track age and disease in peripheral samples from living subjects. As such, this measure may be capturing a cellular program that is activated in senescence. In moving forward, it will be important to evaluate this measure using data from clinical trials, relate it to existing senescence biomarkers, and further evaluate reliability using diverse cell types and inducers. Finally, further instigating the epigenetic changes that accompany senescence may provide important insight into the characteristics of this aging phenotype, facilitating or ability to develop reliable therapeutics to target these cells.

## METHODS

Two datasets were used for training DNAmSen. For both studies, DNAm levels were assessed using the Illumina HumanMethylation450 BeadChip. The first was a dataset from Xie et al., 2018, obtained from Gene Expression Omnibus (accession no. GSE91069). Briefly, early passage (EP) BJ fibroblast, were sequentially passaged to induced RS (28 population doublings). Near senescence cells were also considered, which cells passaged for 14 doublings, that while remaining proliferative, exhibit SA-βgal^+^ staining. OIS cells were produced by infecting EP cells with the H-Ras oncoprotein (H-rasV12), which after 10 days activates a senescence phenotype. Additionally, data from immortalized and transformed cells were also available in comparison to empty vector (EV). For instance, EP cells were infected with either human telomerase catalytic subunit (hTERT) alone, or hTERT and subsequently infected with simian virus 40 large T antigen (SV40) to promote immortalization. To fully transform them, cells infected with hTERT and SV40 were further infected with a H-rasV12 retrovirus, generating a fully immortalized/transformed lineage. Lastly, near senescence cells (mentioned above) were also immortalized via SV40 and/or hTERT, yet although they were infected with H-rasV12 and exhibited increased expression, these cells appeared to resist immortalization.

The second dataset (GSE37066) was from Kock et al., 2013 and Shao et al., 2013 and included DNAm data from MSC characterized as EP, late passage/RS, irradiated, immortalized, or cells reprogrammed into induced pluripotent stem cells (iPSC). As with the data from Xie et al., cells were passaged in culture, and in this case, we considered 3 samples as late passage (12-16 passages). EP was defined as those with only two to three passages in culture. As with the data from Xie et al. immortalization was achieved via overexpression of TERT or TERT and SV40. IRIS was achieved seven days after irradiating cells with 15 Gy. Finally, data was also available from embryonic stem cells (ESC) and iPSC based on reprogrammed MSC.

Elastic net, fitting a binomial logistic regression was used to train a classifier of senescence versus non-senescence. The senescence category included all three types of inducers—RS (n=3 samples from Fibroblasts and n=3 samples from MSC), OIS (n=3 samples from Fibroblasts), and IRIS (n=3 samples from MSC)—totaling 12 senescence samples. These were contrasted against n=10 EP samples (5 from MSC and 5 from Fibroblasts). Briefly, 10-fold cross-validation was used to select a lambda penalty for the elastic net regression, resulting in a classifier based on DNAm levels from 88 CpGs.

Four datasets were used to validate the DNAmSen measure using in vivo samples—GSE40279, GSE51954, GSE42865, GSE63704, GSE56515. In each dataset, the equation resulting from the elastic net from cell cultures (88 CpGs) was applied to estimate DNAmSen. For the datasets from whole blood, epidermis/dermis, and fetal tissue, we tested the association between DNAmSen and chronological age using biweight midcorrelation. For the data from progeroid patients versus controls, and diseased versus healthy lung samples we used step-wise ordinary least squares (OLS) regression to test whether groups differed in their predicted levels of DNAmSen. In model 1, no confounders were considered in the models. However, for model 2, we adjusted fro a surrogate age measure—Horvath epigenetic clock. The epigenetic clock was estimated in accordance with the equation proposed by Horvath^20^ that involves a weighted sum of 353 CpGs.

